# Basement membrane turnover drives filopodial protease-independent invasion

**DOI:** 10.64898/2026.01.05.697631

**Authors:** David Hernandez-Aristizabal, Anotida Madzvamuse, Frédéric Luton, Rachele Allena

**Affiliations:** Université Côte d’Azur, LJAD UMR CNRS 7351, Nice, France; The University of British Columbia, Mathematics Department, 1984 Mathematics Road, Vancouver, V6T 1Z2, Canada; Department of Mathematics and Computational Sciences, University of Zimbabwe, Mt Pleasant, Harare, Zimbabwe; Department of Mathematics and Applied Mathematics, University of Pretoria, Pretoria, 0132, South Africa; Applied Mathematics, University of Johannesburg, PO Box 524 Auckland Park Johannesburg, 2006, South Africa; Université Côte d’Azur, IPMC UMR CNRS 7275, INSERM U1323, Valbonne, France; Institut Universitaire de France, Paris, France

**Keywords:** basement membrane, collagen IV turnover, filopodia, cancer invasion, geometric-surface PDE modelling, evolving surface finite element method, pore

## Abstract

The basement membrane is a specialised nanoporous extracellular matrix mainly composed of collagen IV fibres and laminins. As collagen IV fibres form covalent cross-links, cells cannot freely cross it. Basement membrane breaching marks the transition from in situ to invasive carcinoma, commonly attributed to protease activity. Yet, recent data show that tumour cells can cross basement membranes through protease-independent mechanisms. Experimental evidence suggests that filopodia can play an active role in pore enlargement in extracellular matrices by generating plastic mechanical deformation. Moreover, plasticity can be regulated by covalent cross-link concentration, which we hypothesise can be responsive to collagen IV turnover, thus generating transient weak spots. To test this, we developed a quantitative, biophysical mathematical model describing the interaction between a tumour-cell cluster and a basement membrane using relevant biological assumptions and parameters. The cluster is represented as an evolving, energetic surface and the basement membrane is described as a set of points representing active and inactive links. Simulations show that synchronisation of collagen IV turnover coupled with filopodium extension drives pore enlargement, providing a mechanistic basis for protease-independent invasion. Consistent with experiments, simulations identify two complementary filopodium groups: one driving global degradation and the other promoting local pore enlargement.

## 1. Introduction

The basement membrane (BM) is a specialised extracellular matrix (ECM) that surrounds most tissues [1]. The BM is primarily composed of laminin, collagen IV fibres, perlecan, nidogen, and other associated proteins. Laminins form a non-covalent network that interacts with cell-surface receptors [2], and collagen IV fibres assemble into a covalently cross-linked network. Nidogen and perlecan connect these two networks, thereby stabilising BM integrity [1]. Covalent cross-linking of collagen IV provides mechanical integrity [3, 4], making the BM stiffer than the underlying cells, which highlights their role in bearing mechanical stress in tissues. In addition, the BM functions as a signalling platform and an adhesive substrate for several cellular processes, such as migration and polarisation [1]. Importantly, the BM acts as a physical barrier limiting cellular passage between tissue compartments. Typical BM pores range from 9 to 112 nm [5, 6], whereas cells cannot traverse pores smaller than about 30% of their nuclear diameter (1–3 µm) [7]. This barrier function is critical in processes such as cancer metastasis, where tumour cells must breach the BM [1].

It was long believed that tumour cells required matrix metalloproteinases (MMPs) to traverse the BM [1]. However, clinical trials targeting MMPs failed to reduce cancer mortality [8], and invasion has been observed even in their absence [9], pointing to protease-independent mechanisms. Several experimental observations point to the involvement of cell cytoskeletal structures. Rho/ROCK-mediated contractility regulates BM remodelling during development [10], and actomyosin contractility together with protease activity drives BM perforation during branching morphogenesis [11]. Cancer cells also form filopodium-like protrusions enriched in integrins that initiate adhesion plaques and promote metastatic outgrowth [12]. Filopodia contribute to substrate tethering, environment sensing, and invasion [13–15]. Moreover, actin polymerisation can generate mechanical forces sufficiently large to deform the BM just before invasion [16]. Corinus et al. [17] demonstrated that invasive mammary spheroids extend long filopodia coinciding with invasion sites, and Wisdom et al. [18] showed that filopodia can enlarge pores in ECMs exhibiting mechanical plasticity. Here, plasticity refers to the matrix ability to undergo permanent deformation induced by mechanical stress and microstructural rearrangement [4, 18, 19]. Further, Wisdom et al. [19] showed that the plasticity of reconstituted BMs can be regulated by modulating the density of collagen IV covalent cross-links (CCLs) by varying transglutaminase content. This suggests that not all collagen IV fibres in the BM are covalently linked.

Far from being a static structure, the BM undergoes continuous turnover [20]. In *C. elegans*, between 25% and 48% of collagen IV fibres are replaced within 2–5.5h [21–23]; in *Drosophila*, halftime turnover occurs within 4–14h [24]; and in mice, replacement ranges from 9% after 10 minutes [25] to 50% after 3.5h [26]. From these data, the turnover halftime of collagen IV fibres can be estimated to be between 1 and 12h.

Based on these observations, we hypothesise that CCLs are constantly replaced by ongoing collagen IV turnover, leaving a brief window of opportunity for filopodia to slip into. During turnover, the network may become locally weakened, allowing cancer cells to extend protrusions such as filopodia. These protrusions could in turn disrupt local matrix renewal, eventually leading to pore enlargement. This possible behaviour aligns with the concept of opportunistic BM breaching described in [27], in which immune cells exploit pre-existing weak spots to traverse the BM. Here, we propose that turnover-driven weak spots can be similarly exploited by cancer cells. Moreover, after filopodial retraction, local BM damage may require repair before new collagen IV fibres can be incorporated, thereby delaying CCL replacement. Repair dynamics are likely tied to the synthesis of BM core components: for example, the expression of BM proteins is reduced at invasive regions in colorectal cancer [28], and their downregulation is associated with invasion in mammary cell spheroids [17, 29]. Even in healthy situations, protrusions can help maintain BM perforations, as observed during branching morphogenesis, where such perforations facilitate rapid local expansion [30].

Experimentally testing this hypothesis is challenging, since it would require simultaneous tracking of filopodial dynamics and protein turnover *in situ*. As an alternative, in this work we propose an *in silico*, biophysical model to investigate the conditions under which turnover may enable protease-independent invasion. This requires modelling a tumour-cell cluster (TCC) interacting with a BM containing dynamic collagen IV fibres.

Various computational strategies have been used to model cell–BM interactions. Agent-based models [31–34] capture large-scale tumour evolution but cannot resolve subcellular processes such as filopodium extension. Cells are deformable when forming filopodia, whereas agent-based models fail to describe cell shape morphology and are rule-based. Phase-field models overcome this limitation by implicitly defining the plasma membrane through a continuous field function. They have been successfully applied to model intra- and extra-cellular processes without the need of knowing precisely the location of the plasma membrane [35–38]. However, for describing filopodia, we do need the exact position of the plasma membrane. An alternative is provided by sharp-interface models [39–43], in which the plasma membrane is explicitly treated as the boundary of a continuously deforming domain, ensuring precise membrane localisation. We refer the interested reader to [44] for a detailed comparison between these two approaches.

Among sharp-interface models, the geometric surface approach describes the plasma membrane as an energetic manifold [45, 46] that minimises the Helfrich functional, which describes the elastic energy of biological membranes [47, 48]. This approach has two main advantages: it allows high cell deformation and reduces computational cost, since the plasma membrane is treated as a one-dimensional curve embedded in 2D or a two-dimensional surface embedded in 3D. Furthermore, it can be naturally extended to take into account intracellular processes through the bulk-surface partial differential equations [41, 42, 45, 49–52]. A simplified strategy has been presented in [19, 53] for the interaction between the plasma membrane and ECMs in confined migration. Wisdom et al. [19] and Ju et al. [53] also restrict their description to the cell surface but use simpler mechanical rules: the membrane is modelled as a closed chain of elastic links with preferred lengths and angles following Langevin equations. Though not curvature-based, these models capture local deformation, steric effects, and ECM plastic rearrangements.

Building on the geometric surface PDE (GS-PDE) approach, we developed a biophysical model that captures the interplay between a deformable TCC and a dynamic BM undergoing collagen IV turnover. The TCC is described by an evolving, energetic membrane governed by a GS-PDE derived from the Helfrich functional. This formulation enables large deformations and subsequently the emergence of plasma-membrane protrusions such as long and thin filopodia. The BM is modelled as a fixed barrier discretised by a set of points representing collagen IV CCLs. Each point switches between active and inactive states following a Langevian process to simulate ongoing collagen turnover and exert a repulsive force against the TCC during active states. Such a GS-PDE modelling framework allows us to explore how filopodial activity and collagen IV turnover may together generate conditions favourable to protease-independent invasion. Moreover, the approach is flexible and can be extended to incorporate protease-dependent mechanisms by allowing local matrix degradation in response to biochemical cues [54–56].

The remainder of the paper is organised as follows. In Section 2, we describe the *in vitro* configuration that we aim to reproduce, together with the biophysical modelling framework, parameters of interest, and numerical schemes. Section 3 presents the simulation results, and Section 4 concludes with a discussion comparing our findings with previous experimental observations.

## 2. Materials & Methods

### 2.1. Cells, reagents, confocal immunofluorescence and time-lapse imaging

The MCF10 PSD4/EFA6B knock-out cells were previously described [29]. Cells were grown in DMEM/F-12 (1: 1), horse serum 5%, non-essential amino acids 1%, insulin 10 µg*/*ml, hydrocortisone 1 µg*/*ml, EGF 10 ng*/*ml, cholera toxin 100 ng*/*ml and penicillin (100u*/*ml)-streptomycin (100 µg*/*ml) invasive mammary cell reagents were from Invitrogen (Fisher Scientific, France). 3D culture was performed using Matrigel (5 mg*/*ml) or rat tail Collagen I (2 mg*/*ml; Corning®, Fisher Scientific, France). The collagen I solution was neutralized using 1N NaOH and diluted in PBS to a final concentration of 2 mg/ml. 5 µl of 0,5×106 cells was mixed in a 25 µl drop of Matrigel deposited on a glass coverslip in 24-well plate, or in 150 µl of Collagen I placed in a well of an 8-well Nunc™ Labtek™ dish. For confocal immunofluorescence, after three washes in PBS, the spheroids embedded in Matrigel were fixed in 4% paraformaldehyde for 30 min, washed 3 times and then incubated in PBS containing 2% fish skin gelatin. Then the spheroids were incubated with primary antibodies over-night and counterstained with appropriate fluorescent secondary antibodies, DAPI and phalloidin (Molecular probes, Invitrogen) as indicated in the figure legend. Images acquisition was performed with a confocal microscope TCS SP8 with a HCX PL APO 63X/1.4 objective (Leica Microsystems). The anti-collagen IV antibody was from Millipore (reference AB769). For time-lapse imaging, to form spheroids 50×103 cells were incubated in complete medium for 48h in poly-HEMA coated 35 mm dishes. Size homogenous spheroids isolated by differential low-speed centrifugation were resuspended in 300 µl of fibrillary collagen I (2.5 mg/ml) and plated in 24-well plates. 48h later images were acquired using a Cytation™ 5 imaging multi-mode reader (BioTek Instruments) in a controlled environment. Time-lapse images were captured every 30 min and up to 72h with a 20X/0.45 phase contrast objective using the Gen5 v3.13 software (Biotek Instruments).

### 2.2. Biophysical model

In this section, we develop a biophysical model based on a recently developed GS-PDE approach to describe the interaction between tumour cells and the BM. It is composed of a system of GS-PDEs describing the spatio-temporal evolution of the TCC and a system of ordinary differential equations (ODEs) describing the temporal dynamics of the collagen IV on the BM. This novel GS-PDE modelling approach allows for studying cell invasion in the absence of biochemical degradation of the BM, and for large plasma-membrane deformations.

Before describing the details of the biophysical model, let us consider the *in vitro* configuration that we aim to reproduce, as reported in [17]. Fig. 1**(a)** presents a spheroid made of invasive human mammary cells (nuclei in blue and actin filaments in red) surrounded by a BM (green). This configuration is obtained after 96h of growth in Matrigel and corresponds to the initial state (*t* = 0) of both the subsequent experimental observations and our simulations. Fig. 1**(b)** illustrates a typical invasion sequence, where the spheroid employs both filopodial protrusions and protease degradation to breach the BM (see original Movie S1). Initially, the BM is intact and no filopodium is observable. After 10.5h, long filopodia (up to 20 µm) protrude beyond the BM, and at 33h cells start crossing through enlarged pores—giving rise to the initiation of the invasion.

**Figure 1.**
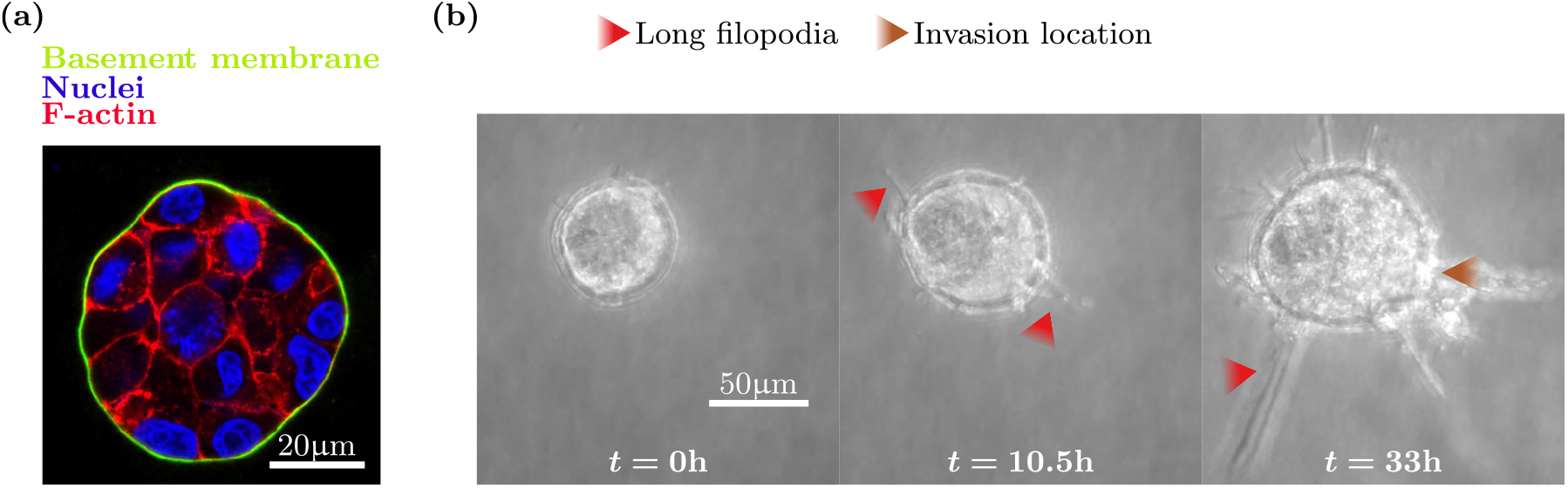
Invasive mammary cell spheroid undergoing BM invasion. **(a)** Representative confocal immunofluorescence image of a MCF10 EFA6B knock-out spheroid grown in Matrigel for 96h and stained for collagen IV (green) to identify the basement membrane, actin filaments (red) and cell nuclei (blue). **(b)** Fixed phase contrast images from a representative movie of a MCF10 EFA6B knock-out spheroid invading the collagen I gel. Scale bar 50µm. The BM is breached by filopodial activity and protease degradation *in vitro* over a time interval of 33h. Long filopodia (up to 70µm) are indicated by red arrowheads, and invasion sites are indicated by a brown arrowhead. See original Movie S1.

To reproduce the behaviour of the spheroid, we developed a GS-PDE modelling framework that couples the evolution of a protruding TCC with the dynamics of the BM. A summary of the resulting framework in its dimensionless form is presented in Fig. 2 and its derivation is given in the subsequent subsections.

**Figure 2.**
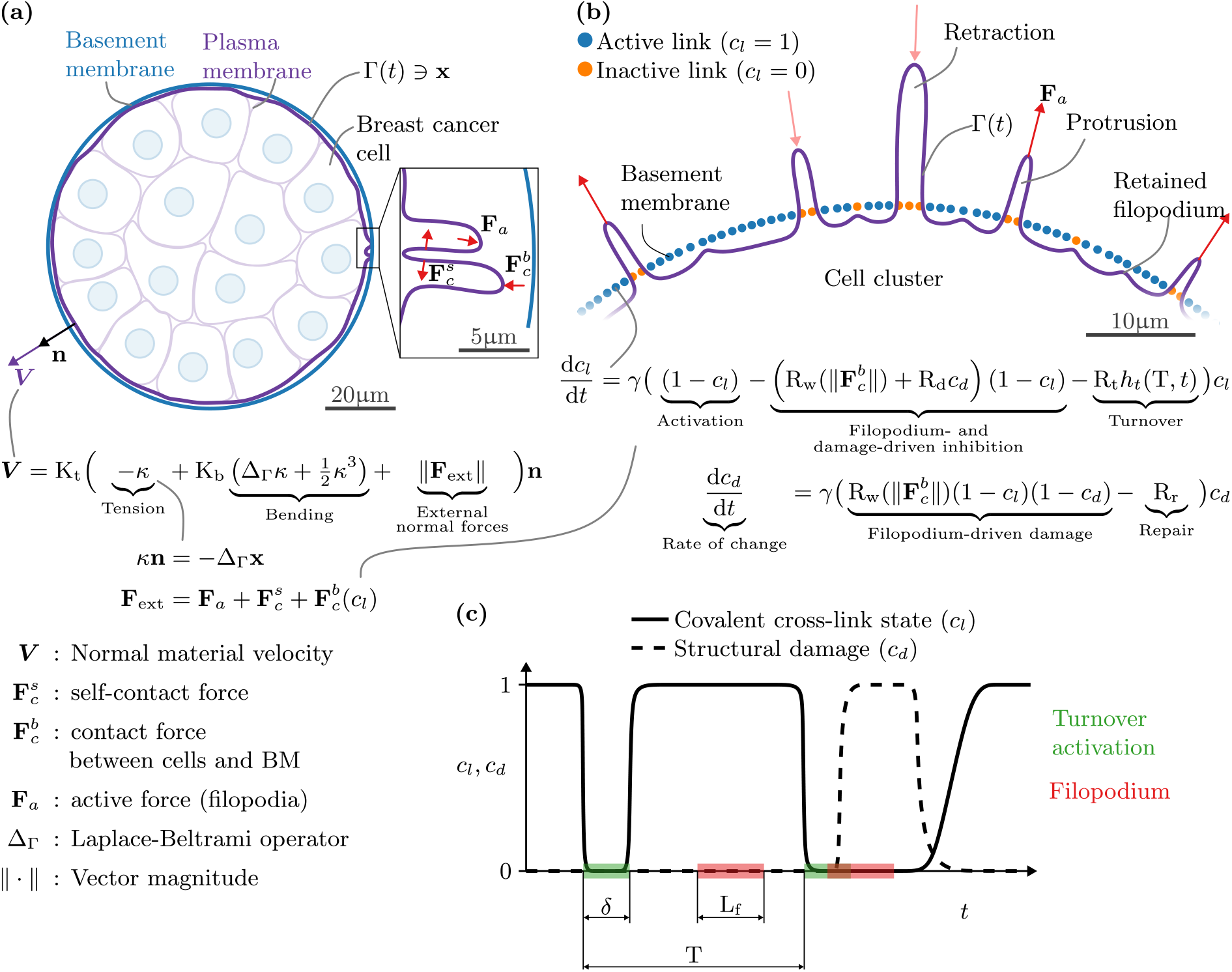
Dimensionless GS-PDE modelling framework of TCC interacting with the BM. The system of equations presented corresponds to the dimensionless form Eqs. (14) to (17). **(a)** A TCC, corresponding to a spheroid of invasive mammary cells. The TCC is described by an evolving surface Γ(*t*) (purple line) interacting with the BM (blue line). Its material velocity ***V*** is governed by a 4th-order GS-PDE that minimises the Helfrich functional, which accounts for curvature (*κ*)-dependent elastic energy. It balances both internal and external normal forces, including filopodial activity (**F**_*a*_), self-contact (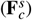), and contact with the BM (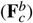). **(b)** Filopodia slip into gaps in the BM accessible during collagen IV turnover. The BM is represented as a set of CCLs, depicted as points that can be either active (blue) or inactive (orange). Each CCL is governed by two ODEs: one for its activity state (*c*_*l*_)—it switches between active (1) and inactive (0) at a frequency of 1*/*T due to turnover—and one for reversible structural damage (*c*_*d*_)—it grows if filopodia act on inactive CCLs. Reactivation of a CCL is inhibited by filopodium presence (R_w_(**F**_*a*_)) or pre-existing damage. **(c)** An illustrative example of the temporal dynamics of the previous system of ODEs. The continuous line shows CCL state and the dashed line shows the BM structural damage in time. The green bands indicate turnover events, at every T for a time interval of *δ*, and the red bands indicate filopodium formation with lifespan L_f_.

#### 2.2.1. The GS-PDE model formulation for the evolution of the tumour-cell cluster

As shown in Fig. 2**(a)**, we describe the TCC by a closed curve (purple line), corresponding to the part of the plasma membrane of cells interacting with the BM (blue line). Then, let us consider an evolving, smooth, orientable, energetic, closed curve Γ(*t*) ∋ **x**(*t*) in ℝ^2^, for *t*∈ [0, *t* _*f*_], with *t* denoting time and *t* _*f*_ final time. Its evolution law (written in Fig. 2**(a)**) arises as a force balance equation posed at each material point on the evolving surface in the outward-pointing normal direction **n**. This approach easily generalises straight forwardly to surfaces embedded in ℝ^3^ [57]. For the derivation, let us assume that the internal energy of the curve is dependent on the geometry and is given by the Helfrich functional [47, 57, 58]:

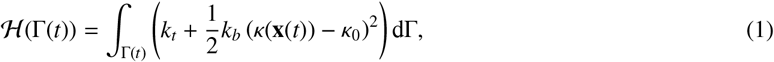

where *k*_*t*_ and *k*_*b*_ are positive parameters associated with the surface tension and bending, respectively. *κ* refers to the mean curvature given by the identity [58]:

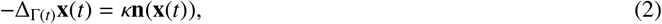

where Δ_Γ(*t*)_ is the Laplace-Beltrami operator [57], and *κ*_0_ is a spontaneous mean curvature.

Assuming that the curve tends to minimise its energy, its evolution is given by the first variation of the internal energy in the direction of **n** plus the action of external normal forces [47, 57]:

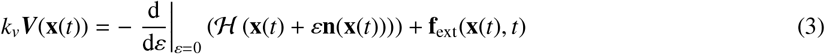

where ***V*** and **f**_ext_ refer respectively to the material velocity and physically relevant external forces in the normal direction. *k*_*v*_ is a positive parameter representing viscosity. This leads to the following nonlinear GS-PDE governing the evolution of Γ(*t*) [45, 47, 58]:

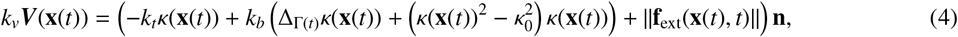

where ||**f**_ext_|| is the magnitude of the external normal forces.

We decompose external normal forces as:

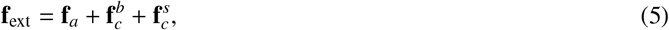

where **f**_*a*_ represents active normal forces generated by actin filaments acting on the cell membrane, and 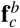 and 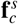 represent contact normal forces between the TCC and the BM and the TCC with itself, respectively. See Fig. 2**(a)** for an illustration of these forces.

In this work, the active force leads the formation of filopodia. Each filopodium is represented by a localised force present during a given period of time *l* _*f*_ and applied on a length *w*_*f*_ . In other words, we consider that a filopodium *i* is generated by pushing the membrane on 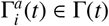 at the time interval 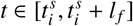. 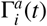 is a connected arc of size *w*_*f*_, and 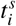 refers to the activation time. For a given number of filopodia *N*_*f*_ appearing in the time interval of interest [0, *t* _*f*_], we can define 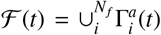 as a subset of Γ(*t*) where filopodia are active at time *t*. Therefore, the active force can be written as:

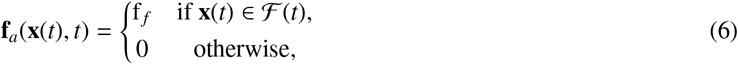

where f _*f*_ is the magnitude of the filopodium force. We randomly define both the position of 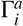 and 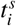. Lastly, *N*_*f*_ is calculated from the filopodium density *ρ*_*f*_, that indicates the fraction of filopodial length over the curve, as:

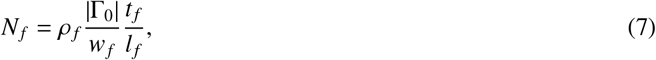

where | · | is the arc length.

We proceed to model contact forces, these are computed as distance functions similarly to [45]. The closer two points are to one another, the larger the force is. The contact force with respect to the BM is defined as:

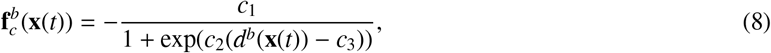

where *c*_1_, *c*_2_ and *c*_3_ are positive parameters, and *d*^*b*^(**x**(*t*)) denotes the minimum distance from **x**(*t*) ∈ Γ(*t*) to the BM.

In the case of self-contact, Eq. (8) is extended to exclude interactions between adjacent points and to determine the force sign as follows:

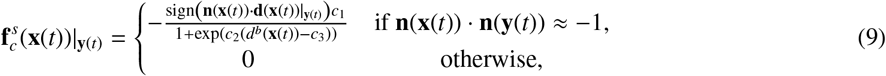

where 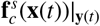 is the self-contact force on **x**(*t*) ∈ Γ(*t*) exerted by **y**(*t*) ∈ Γ(*t*), and **d**(**x**(*t*))|_**y**(*t*)_ is the distance vector from **x**(*t*) to **y**(*t*). An illustration of the different scenarios provided by Eq. (9) is shown in Fig. 3.

**Figure 3.**
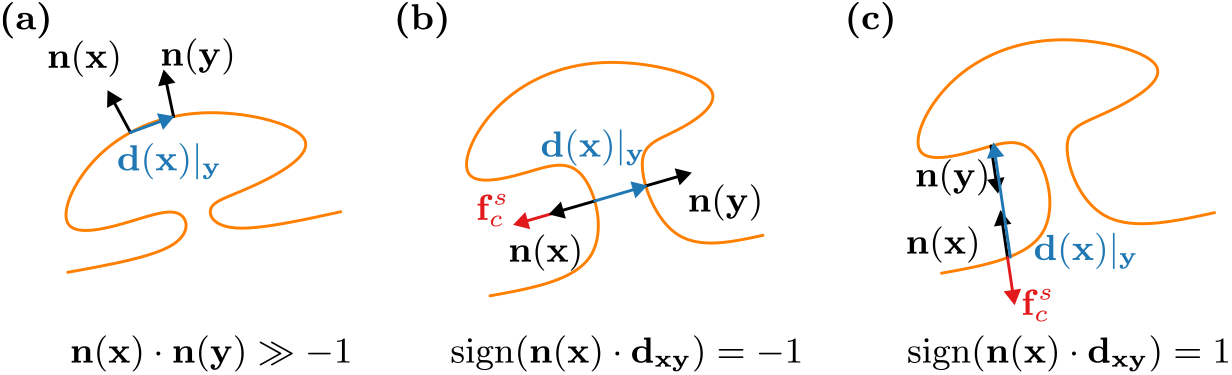
Computation of self-contact forces. Following Eq. (9), three scenarios are illustrated: **(a)** two adjacent points close to each other do not exert a contact force, **(b)** two non-adjacent points exert contact forces directed along the normal vector, and **(c)** two non-adjacent points exert contact forces directed opposite to the normal vector.

#### 2.2.2. The temporal dynamics of the basement membrane turnover

In the BM, collagen IV fibres are joined by CCLs [3]. Consequently, tumour cells cannot cross the BM as they cannot break such links. However, collagen IV fibres undergo turnover with half-life times estimated between 1 and 12h [21–26]. During a fibre’s turnover process, the CCL is temporally lost.

We shall model the BM as a set of points corresponding to CCLs between collagen IV fibres (Fig. 2**(b)**). These points are distributed at equal intervals to recreate a BM porosity, with a mean pore size of 50 nm [5, 6]. Each point can be in an active state (blue points in Fig. 2**(b)**), indicating the presence of a CCL, or in an inactive state (orange points in Fig. 2**(b)**), indicating its absence. Active CCLs exert a contact force (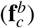) against the TCC, thereby blocking filopodia formation and extension, while inactive ones offer no resistance, allowing filopodia to penetrate across the BM.

Our main hypothesis is that filopodia exploit local weakening of the BM during the replacement time to occupy the space of collagen IV fibres. Yet, this replacement time is generally short, given the abundance of collagen IV fibres in the environment. Therefore, only filopodia already pushing the BM at the moment of turnover are likely to occupy this space. Eventually, such filopodia retract, leaving an open space that new collagen IV can reoccupy.

We further assume that this temporal invasion causes reversible damage to the matrix structure that need repair before the stabilisation of a new collagen IV fibre. Therefore, this damage delays the reactivation of the CCL after filopodium retraction. This can be seen as a loss of affinity due to the rearrangement of the BM structure around crossing filopodia. Biologically, the repair speed can be related to the synthesis of BM core components, whose expression has been reported to be reduced at invasive regions in colorectal cancer progression [28] and in invasive mammary cell spheroids [17, 29].

Based on these experimentally-observed ideas, we describe the state of each BM point using two ODEs: one for the activity of CCLs, *c*_*l*_, and one for structural damage on the BM, *c*_*d*_. Let us consider 0 ≤ *c*_*l*_, *c*_*d*_ ≤ 1, where *c*_*l*_ = 0 corresponds to an inactive CCL and *c*_*l*_ = 1 to an active one, while *c*_*d*_ = 0 represents undamaged structure and *c*_*d*_ = 1 damaged structure. In the absence of filopodium forces and turnover, *c*_*l*_ = 1 and *c*_*d*_ = 0 should be stable equilibrium points. In contrast, sustained filopodium contact forces should shift the stability towards *c*_*l*_ = 0 and *c*_*d*_ = 1, representing local disruption and damage of the BM. This behaviour is captured by the following temporal system of ODEs:

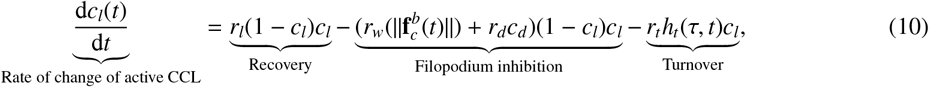

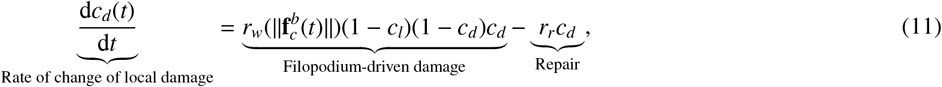

where *r*_*l*_, *r*_*w*_, *r*_*d*_, *r*_*t*_ and *r*_*r*_ respectively denote the rates of CCL replacement, replacement inhibition due to filopodium force, replacement inhibition due to damage, degradation due to turnover, and repair rate after filopodium retraction, respectively. As can be seen, *r*_*w*_ is a dynamic term depending on filopodium contact force on the BM 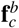. *r*_*w*_ can be defined as a saturating Hill function:

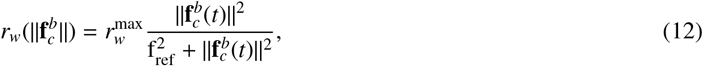

Where 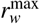 is the maximum rate of inhibition and f_ref_ is a reference force scale. Moreover, in the turnover term, the function *h*_*t*_ is a periodic function with a period of *τ* that activates turnover for a time interval of *δ* (see Fig. 2**(c)**).

The temporal dynamics of the CCLs are illustrated in Fig. 2**(c)**. Initially, the CCL is active and undamaged. During a turnover event (green zones), it becomes inactive. In the absence of filopodia, the CCL reactivates and the BM remains undamaged. By contrast, if a filopodium (red zones) coincides with turnover, reactivation is inhibited and the BM structure is damaged. After filopodium retraction, the structure first repairs before CCL activity is restored. Thus, persistent inactive states of CCLs depend on the temporal coincidence of filopodia with collagen IV turnover events.

#### 2.2.3. Dimensionless model

The interaction between the TCC and the dynamic BM is governed by Eqs. (2), (4), (10) and (11) with the parameters listed in Table 1. Let us now consider the following dimensionless 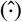 quantities:

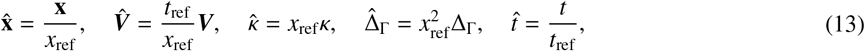

where *x*_ref_ and *t*_ref_ are reference length and time, respectively. We shall define the reference length as the radius of a tumour cell (8 µm). This allows us to interpret *k*_*t*_*/x*_ref_ as a reference membrane tension and 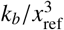 as a reference membrane bending stress, see Table 1. The reference time *t*_ref_ is defined as the time required for a filopodium to extend a distance equal to the radius of tumour cells, 4 min (at rate 2 µm*/*min) [59]. Hence, the term *k*_*v*_ *x*_ref_*/t*_ref_ can be interpreted as the reference viscous force associated with moving at speed *x*_ref_*/t*_ref_.

**Table 1:**
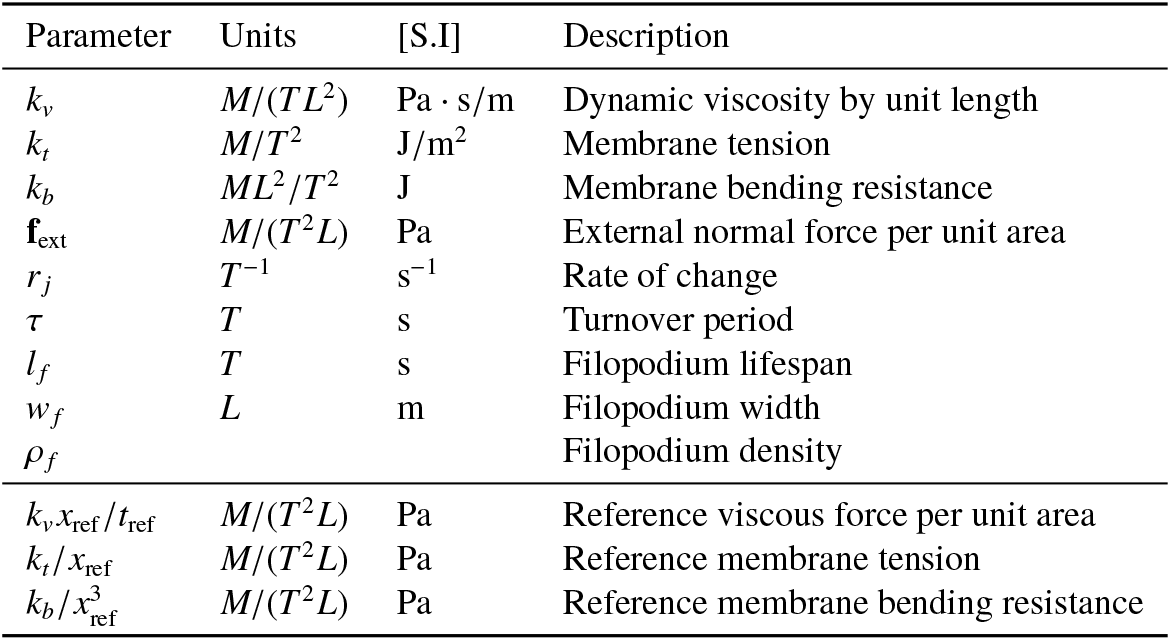
Physical parameters of the GS-PDE model and derived quantities. *M, L* and *T* refer respectively to mass, length and time. [S.I.] refers to international system of units. *r* _*j*_ refers to *r*_*l*_, *r*_*w*_, *r*_*d*_, *r*_*t*_ and *r*_*r*_.

| Parameter | Units | [S.I.] | Description |
| --- | --- | --- | --- |
| $k_v$ | $M/(TL^2)$ | $\text{Pa} \cdot \text{s/m}$ | Dynamic viscosity by unit length |
| $k_t$ | $M/T^2$ | $\text{J/m}^2$ | Membrane tension |
| $k_b$ | $ML^2/T^2$ | $\text{J}$ | Membrane bending resistance |
| $\mathbf{f}_{\text{ext}}$ | $M/(T^2L)$ | $\text{Pa}$ | External normal force per unit area |
| $r_j$ | $T^{-1}$ | $\text{s}^{-1}$ | Rate of change |
| $\tau$ | $T$ | $\text{s}$ | Turnover period |
| $l_f$ | $T$ | $\text{s}$ | Filopodium lifespan |
| $w_f$ | $L$ | $\text{m}$ | Filopodium width |
| $\rho_f$ | | | Filopodium density |
| $k_v x_{\text{ref}}/t_{\text{ref}}$ | $M/(T^2L)$ | $\text{Pa}$ | Reference viscous force per unit area |
| $k_t/x_{\text{ref}}$ | $M/(T^2L)$ | $\text{Pa}$ | Reference membrane tension |
| $k_b/x_{\text{ref}}^3$ | $M/(T^2L)$ | $\text{Pa}$ | Reference membrane bending resistance |

Replacing Eq. (13) into Eqs. (2), (4), (10) and (11) and dropping the hat notation for the sake of simplicity yields the following dimensionless system of equations:

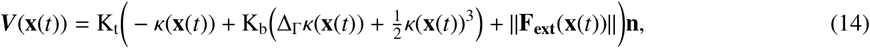

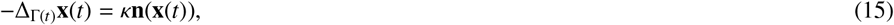

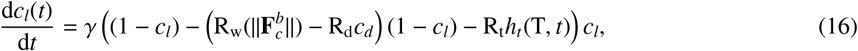

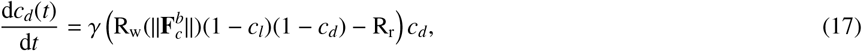

where K_t_, K_b_, *γ*, R_w_, R_d_, R_t_, R_r_ and T are dimensionless parameters, defined in Table 2. K_t_ represents the ratio of membrane tension to viscous forces, while K_b_ represents the ratio of bending stiffness to tension. *γ* and T are the turnover speed and period relative to the time required for the formation of a filopodium of length equal to the radius of tumour cells. R_w_, R_d_, R_t_, R_r_ respectively describe the rates of inhibition of link recovery due to protrusive forces, damage-induced inhibition, link turnover, and damage repair, all with respect to the link recovery rate. The dimensionless external normal forces are given by:

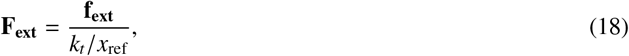

which represents the ratio between external and tension forces. Lastly, the parameters defining filopodium formation are also presented in their dimensionless form in Table 2; W_f_ and L_f_ are respectively normalised filopodium width and lifespan.

**Table 2:** Dimensionless parameters of the GS-PDE modelling framework (Fig. 2). The first part of the table shows the parameters fixed throughout all the simulations. The second part of the table contains the reported ranges of the parameters varied in this study: density, width and lifespan of filopodia and turnover period and repair rate. Reference length: 8 µm (cell radius); reference time: 4 min (time to extend a filopodium to the size of the cell radius). ∗Values were estimated based on the response of the model, while the other values were taken from literature.

| Parameter | Description | Value | Source |
| --- | --- | --- | --- |
| $K_t$ | Ratio between tension and viscous forces | 0.01 | For a filopodium retraction speed equivalent to $2 \mu\text{m}$ [59] |
| $K_b$ | Ratio between bending and tension forces | 0.0001 | Estimated from [60, 61] |
| $F_a$ | Ratio between active and tension forces | | To calibrate for filopodium extension speed equivalent to $2 \mu\text{m}$ [59] |
| $\gamma$ | Normalised turnover speed | 200* | To ensure fast link activation/deactivation |
| $R_w^{\max}$ | Maximum ratio between rates of CCL inhibition by force and recovery | 2* | To ensure recovery inhibition by filopodium presence |
| $R_d$ | Ratio between rates of CCL inhibition by damage and recovery | 2* | To ensure recovery inhibition by damage |
| $R_t^{\max}$ | Maximum ratio between rates of CCL turnover and recovery | 2* | To ensure link deactivation during turnover |
| $\rho_f$ | Density of filopodia | $20 - 50 \frac{\mu\text{m fil}}{100\mu\text{m BM}}$ | [17] |
| $W_f$ | Normalised width of filopodia | 0.0125 – 0.5 | Equivalent to $0.1 - 4 \mu\text{m}$ [17, 59] |
| $L_f$ | Normalised lifespan of filopodia | 7.5 – 90 | Equivalent to 30 – 360 min [17] |
| $R_r$ | Ratio between rates of repair and CCL recovery | $0.000625 - 0.01^*$ | To capture fast and slow repair |
| $T$ | Normalised turnover period | 30 – 260 | Equivalent to turnover halftime 1 – 12h [21–26] |

### 2.3. Parameters of the model

The dimensionless parameters of the system are listed in Table 2. The first block refers to parameters fixed throughout all the simulations, either taken from the literature or calibrated to match expected behaviours. We estimated the membrane properties from reported measurements of tension in HeLa cells and bending resistance in red blood cells [60, 61]. Although these are not tumour cells, we assumed that the ratio between tension and bending resistance is comparable. We calibrated the parameters controlling BM dynamics to reproduce the behaviour shown in Fig. 2**(c)**, namely rapid activation/deactivation of CCLs during turnover and efficient inhibition of recovery. Lastly, we defined filopodial and viscous forces to reproduce an average filopodial speed of 2 µm*/*min (= *x*_ref_*/t*_ref_) [59].

The second block lists the parameters varied in this study, with ranges derived from experimental observations. We characterised filopodial activity by density (*ρ*_*f*_), width (W_f_), and lifespan (L_f_), with values reported for invasive and non-invasive mammary cell spheroids [17]. BM stability was determined by the turnover period (T) and the repair rate (R_r_). We estimated turnover ranges from fluorescence recovery after photobleaching experiments [21–26] by taking the inverse of the measured recovery-rate constant. For R_r_, no experimental data are available; hence, we defined a range capturing fast and slow repair regimes.

To analyse the conditions leading to BM invasion, we performed a sensitivity analysis varying the parameters listed in the second block of Table 2. We varied the parameters following a central composite design to construct a response surface [62–64]. For each parameter set, we ran a simulation for an equivalent time of 48h in a small spheroid (radius equal to *x*_ref_). The choice of reducing the spheroid size was done to reduce the computational cost. Nevertheless, the other parameters rested in their biologically relevant range. Thus, this reduced spheroid can be regarded as a sample surface of a real-size spheroid.

We measured two response variables: maximum BM degradation (*ϕ*), measured as the fraction of inactive CCLs over the total number of CCLs, and maximum pore size (*p*_*s*_). For the response surfaces, we fitted a quadratic metamodel of the form:

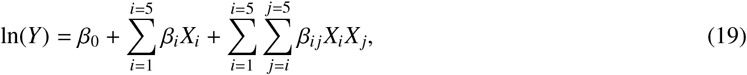

where *X*_*i*_ are the parameters normalised in [ −1, 1] and *Y* is the output response. The responses were logarithmically transformed to improve model accuracy, and some of the parameters were also transformed to better represent their variation. The parameter ranges and normalisation scheme are given in Table 3.

**Table 3:** Parameter space of the sensitivity analysis. Parameters were varied either linearly or quadratically. The linear was used to span biologically reported ranges, while the quadratic one was used to capture variation around a reference value. The columns −1, 0, and 1 indicate extreme and medium values that the normalised variables *X*_*i*_ refer to.

| Parameter | Scale | -1 | 0 | 1 | Observation |
| --- | --- | --- | --- | --- | --- |
| T | Linear | 60 | 145 | 230 | Equivalent to turnover halftime between 2.8 and 10.62h [21–26] |
| $R_r$ | Quadratic | 0.000625 | 0.0025 | 0.01 | For fast and slow repair |
| $\rho_f$ | Linear | 0.22 | 0.36 | 0.5 | Equivalent to density of 1 $\mu\text{m}$-width filopodia in [17], from 22 to 50 filopodia per 100 $\mu\text{m}$ |
| $W_f$ | Quadratic | 0.0625 | 0.125 | 0.25 | Equivalent to half and twice the common reported width 1.0 $\mu\text{m}$ in [17] |
| $L_f$ | Quadratic | 6 | 12 | 24 | Equivalent to half and twice the most frequent filopodium lifespan 48 min in [17] |

### 2.4. Numerical method

The numerical framework comprises the system of Eqs. (14) to (17). Eqs. (14) and (15) describe the motion of the TCC and are solved using the evolving surface finite element method developed in [57, 58, 65–67], which has been applied to cell migration [45] and plant infection [46]. Eqs. (16) and (17) govern the dynamics of the CCLs and are integrated using the backward Euler time-stepping scheme, which is unconditionally stable and therefore allows for larger use of time steps Δ*t* without compromising accuracy. These two systems of equations are coupled through the contact force between the BM and the TCC. The numerical algorithm was implemented using FEniCSx [68–71], and it is available online on Filopodia to pores.

For stability of the GS-PDE, it is necessary to ensure a good node distribution in time [45, 58]. This is achieved by adding a tangential component in the velocity [42, 45, 57, 58, 73]. We remark that this tangential movement does not affect the shape of the curve and hence the force balance is still satisfied.

Here, we compute the tangential displacement using the De Boor’s algorithm adapted to closed curves (see Fig. 4) [42, 68, 72]. Let Γ_*h*_(*t*) be the finite element (FE) polygonal curve (black line in the figure) with node positions 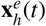 at time *t*. Now, let us assume that the curve undergoes deformation from *t* to *t* + Δ*t*. In general, the nodes of the deformed mesh 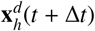 are not regularly distributed. To add an appropriate tangential movement, we construct an interpolating cubic B-spline Γ_*s*_ (red line in the figure) representing a smooth approximation of Γ passing through all the FE nodes. Next, we compute the total arc-length |Γ_*s*_| and calculate the optimal arc-length by element:

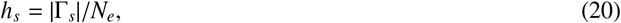

with *N*_*e*_ the number of finite elements. We now want to displace the nodes on Γ_*s*_ such that the arc-length between each node is *h*_*s*_. Since Γ_*s*_ is a closed curve, there are infinite possible configurations satisfying this condition. To define a unique configuration, we select the equidistributed node set that minimises the displacement:

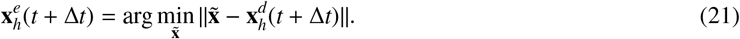

**Figure 4.**
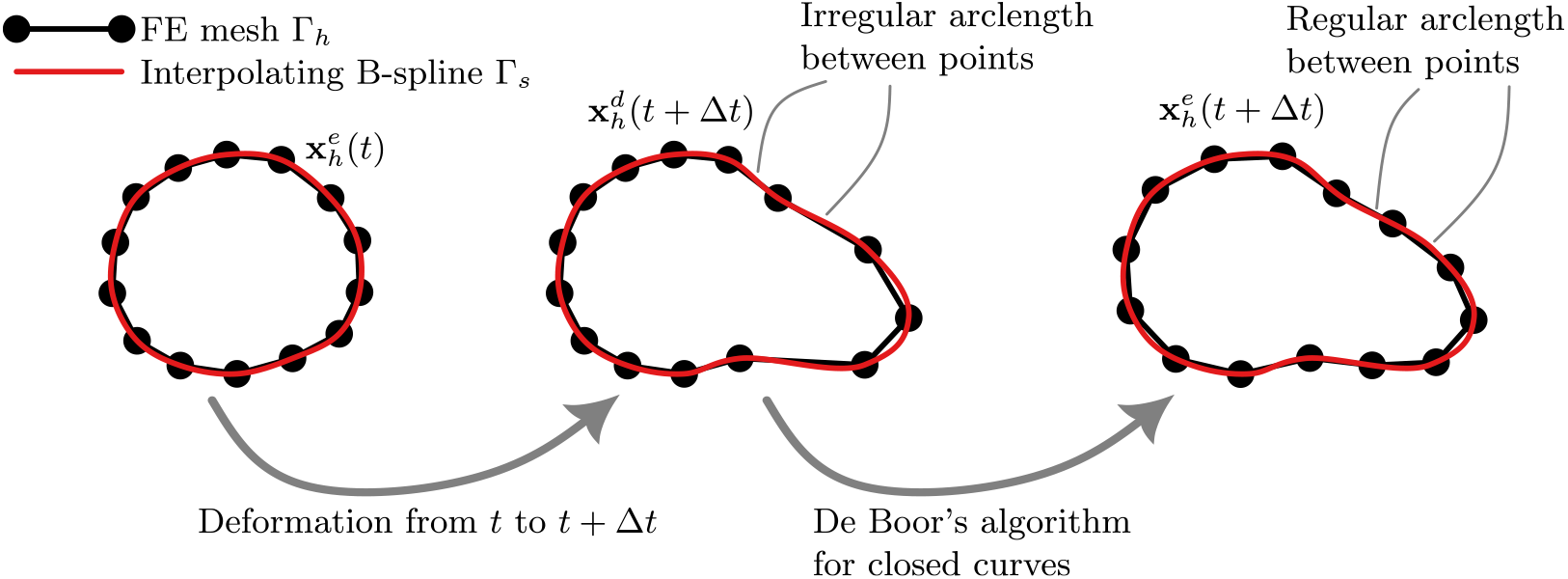
A numerical scheme for mesh equidistribution on curves during deformation. An interpolating cubic B-spline Γ_*s*_ (red line) is constructed from the nodes (black points) of the FE mesh Γ_*h*_ (black line). The nodes are relocated in the spline ensuring a constant arc-length using the De Boor’s algorithm adapted to closed curves [42, 68, 72].

## 3. Results

### 3.1. Convergence

To select an appropriate spatial and temporal discretisation, we performed a convergence study. We considered the extension of a filopodium with W_f_ = 0.125 from a circular cell of radius *x*_ref_. We imposed a velocity of twice the average velocity (2*x*_ref_*/t*_ref_) for a simulation time *t*_ref_, yielding a filopodium length of 2*x*_ref_. Fig. 5**(a)** shows the cell before and after extension.

**Figure 5.**
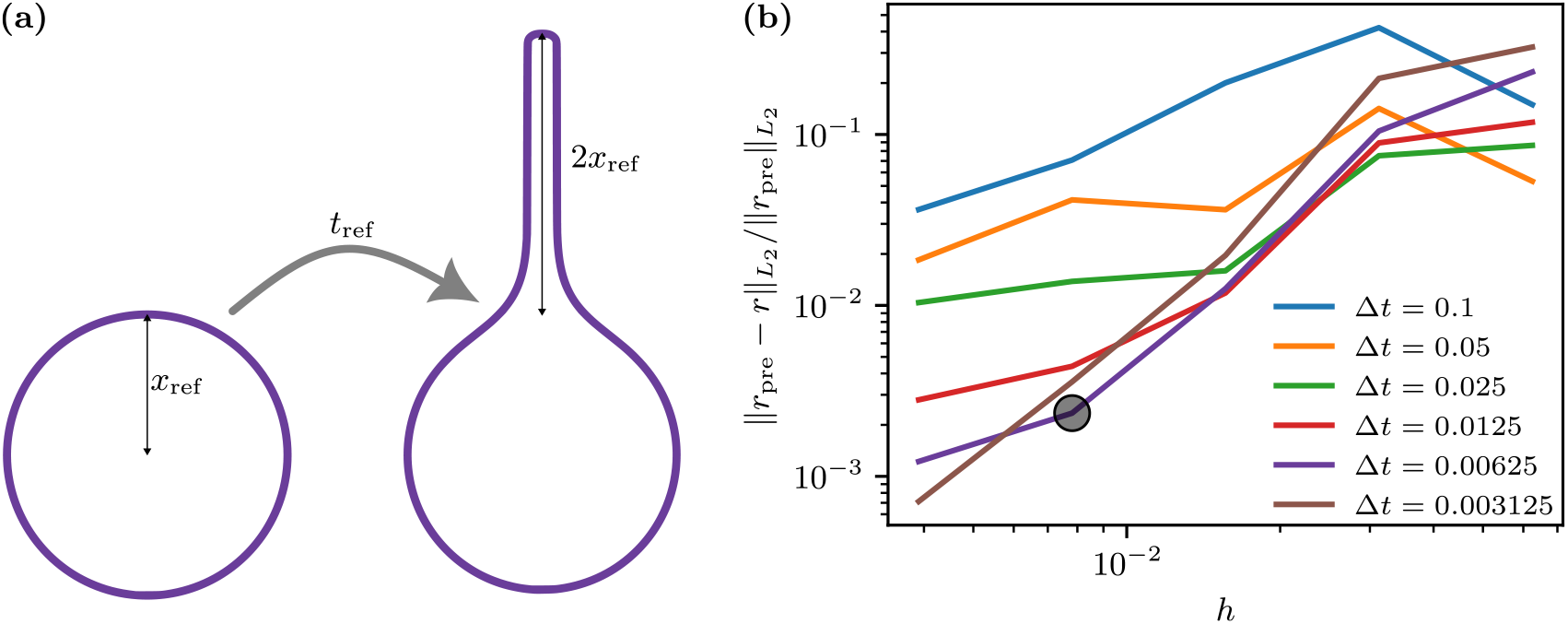
Convergence of the TCC model for filopodium extension at twice the average speed. **(a)** Initial and final configuration of the cell used for convergence measurements. **(b)** Normalised relative error as a function of time step Δ*t* and element size *h. r* denotes radial position of points 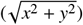, for a given *h* and Δ*t*. The subscript *pre* refers to the previous, finer discretisation with *h/*2 and Δ*t/*2. || · ||_*L*2_ denotes the *L*_2_-norm in Γ × [0, *t* _*f*_] [57]. The black point indicates the chosen parameters for the simulations, where the relative error is about 0.002.

For each discretisation (*h*, Δ*t*), we computed the relative error with respect to a previous finer discretisation (*h/*2, Δ*t/*2) using the parametric function:

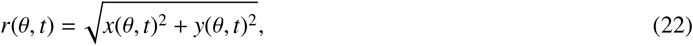

where (*x*(*θ, t*), *y*(*θ, t*)) ∈ Γ(*t*). The relative error was defined as:

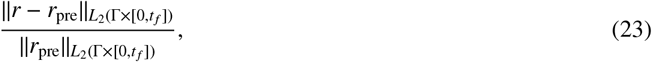

where the subscript *pre* denotes the previous, finer discretisation, and 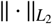 denotes the *L*_2_-norm in Γ × [0, *t* _*f*_] [57].

Fig. 5**(b)** shows the error as a function of *h* and Δ*t*. The error varying both *h* and Δ*t* are shown. We chose *h* = 0.0078125*x*_ref_ and Δ*t* = 0.00625*t*_ref_, corresponding to the black point in the graph. This yields a relative error of about 0.002 and keeps the time step small enough to capture turnover dynamics. For the BM, we used the same time step and a spatial resolution of 0.00625*x*_ref_, equivalent to 50 nm to match the average BM pore size [5, 6]. Due to the discretisation, the smallest time change that can be captured is Δ*t*; hence, we set the turnover interval *δ* = Δ*t*, see green zone in Fig. 2**(c)**.

### 3.2. Collagen-IV-fibre turnover allowing pore enlargement in basement membranes

In this section, we describe a potentially invasive spheroid *in silico* comparing its evolution to the *in vitro* case shown in Fig. 1. We considered as the initial configuration a TCC of diameter 92µm (matching the average diameter of the spheroid *in vitro*) surrounded by an intact BM—all CCLs are active at *t* = 0. We considered that invasion was possible in spheroids with pores reaching a size of at least 2µm [7, 17]. Additionally, we restricted the parameters to the reported values listed in Table 2.

Since no data exist for the repair rate, we first tested a scenario with no damage (*c*_*d*_ = 0), which is equivalent to assuming instantaneous repair. Even under rapid turnover and high filopodial activity, no invasion was observed: pores remained too small, blocking cell passage. Then, we assumed a repair rate of 0.125*/*min (equivalent to R_r_ = 0.0025), which corresponds to ∼ 8 min for BM recovery after filopodium retraction. Under these conditions, we observed potential invasion sites for a turnover period of 4h with 50 filopodia per 100 µm, each 1 µm wide and with a 48-min lifespan, in a spheroid of 92µm diameter.

Fig. 6 shows the obtained protease-independent invasion *in silico*. In the simulation (see Fig. 6**(a)**, and Movies S2 and S3 for a dynamic illustration), a similar sequence to the *in vitro* case (Fig. 1**(b)**) is observed: at 10.5h, long filopodia extend beyond the BM, and at 22h, a pore of 2 µm has formed—which is large enough for invasion [7]. The maximum filopodium length as a function of time is shown in Fig. 6**(b)**. Filopodia longer than 10µm begin to emerge at about 4h, and the maximum length 57.1µm is reached at about 9h. Fig. 6**(c)** compares the pore size distribution *in silico* and *in vitro* [17]. In the former, the distribution at 22h is presented. In the latter, three distributions are shown corresponding to mammary cell spheroids under non-invasive conditions (WT), high filopodial activity (Bleb) and invasive conditions (DICS). As we can see, although most of the pores *in silico* are small (less than 0.25 µm size) as in the non-invasive conditions, the larger pores correspond to invasive spheroids *in vitro*. The corresponding degradation dynamics are shown in Fig. 6**(d)**. BM degradation rapidly reaches a quasi-stationary state at ∼12%, while pore size continues to grow. Initially, this degradation corresponds to small pores (less than 1µm), but over time larger pores (up to 2µm) appear without further increase in overall degradation. This behaviour suggests that BM damage allows synchronisation between turnover and filopodial activity. Since filopodia are randomly distributed, several may act on the same CCL at different times. The damage-induced reactivation delay allows successive filopodia to occupy the same site by preventing its closure. In addition, subsequent turnover events in neighbouring CCLs yield pore enlargement if filopodia are present, leading to progressive widening.

**Figure 6.**
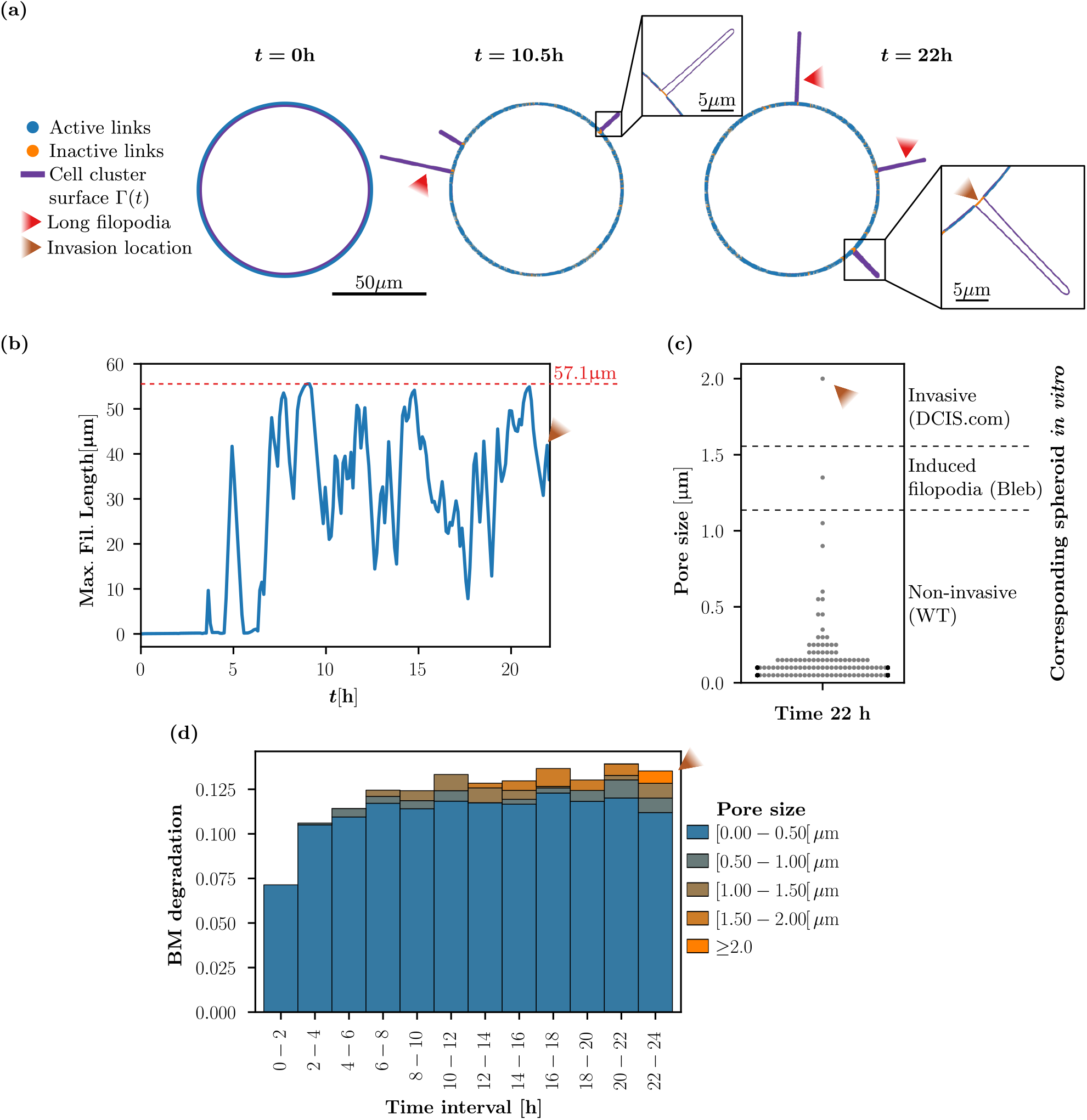
In our GS-PDE modelling framework, collagen IV turnover offers a window of opportunity for BM invasion of a TCC. **(a)** Model simulations reproduce invasive dynamics without protease degradation in a time interval of 22h in a spheroid of 92µm diameter. The simulation used a turnover period of 4h and a repair rate of 0.125min^−1^, considering 50 filopodia per 100µm, each 1µm wide, with a 48-min lifespan. The BM is represented by blue (active) and orange (inactive) points (CCLs), and the TCC is represented by a purple curve (cell plasma membranes Γ(*t*)). The zoomed squares show a pore of 1.3µm at 10.5h and another of 2µm at 22h. **(b)** Maximum filopodium length is bounded. The graph shows the maximum filopodium length as a function of time. The maximum length was 57.1µm (dashed red line). **(c)** Pore size distribution of the simulation at 22h compared to the distribution presented in Fig. 4E in [17]. The dashed lines indicate the limiting maximum pore size (calculated as the mean plus twice the standard deviation) of the different mammary cell spheroid types evaluated in [17]. WT refers to wild-type non-invasive spheroids; Bleb refers to spheroids treated with blebbistatin to induce filopodia; and DCIS refers to invasive spheroids where filopodia are also more abundant than in WT. The largest pore size *in silico* model (brown arrowhead) corresponds to pore sizes seen in invasive spheroids *in vitro*. **(d)** BM degradation reaches an almost stationary state, while pores are locally enlarged. The graph depicts the mean degradation of the BM during time intervals of 2h. The coloured bars indicate degradation grouped by pore size. The dark blue bars represent small pores and the orange portions represent larger pores, scaled from 0 to ≥ 2 µm.

These results indicate that it is plausible that filopodia exploit collagen IV turnover to enlarge BM pores, offering a mechanistic basis for protease-independent invasion.

### 3.3. Sensitivity analysis of basement membrane and filopodial parameters

We next analysed how BM and filopodial parameters influence invasion by performing a sensitivity analysis. The results of the sensitivity analysis are summarised in the response surfaces shown in Fig. 7. These were obtained by fitting metamodels for maximum BM degradation and maximum pore size using Eq. (19). The fitted coefficients are listed in Table 4, where only significant parameters were considered. The metamodels showed high accuracy, with coefficients of determination (*R*^2^) close to 1.

**Table 4:** Metamodels for the maximum BM degradation (*ϕ*) and the maximum pore size (*p*_*s*_). The *R*^2^ column indicate the coefficient of determination of each model. The rest of the columns indicate the coefficient (*β*, see Eq. (19)) for each parameter. The parameters not shown have a coefficient of zero in both metamodels.

| $Y$ | $R^2$ | T | $R_r$ | $\rho_f$ | $W_f$ | $L_f$ | $TR_r$ | $T\rho_f$ | $TW_f$ | $TL_f$ | $R_r\rho_f$ | $R_rW_f$ | $R_rL_f$ | $T^2$ | $R_r^2$ |
| --- | --- | --- | --- | --- | --- | --- | --- | --- | --- | --- | --- | --- | --- | --- | --- |
| $\ln(\phi)$ | 0.98 | -0.6 | -0.8 | 0.4 | -0.3 | 0.1 | 0.0 | 0.0 | 0.0 | -0.1 | -0.1 | 0.1 | 0.3 | 0.4 | 0.3 |
| $\ln(p_s)$ | 0.93 | -1.0 | -0.8 | 0.5 | -0.4 | 0.2 | 0.3 | -0.2 | 0.2 | 0.0 | -0.2 | 0.3 | 0.3 | 0.9 | 0.0 |

**Figure 7.**
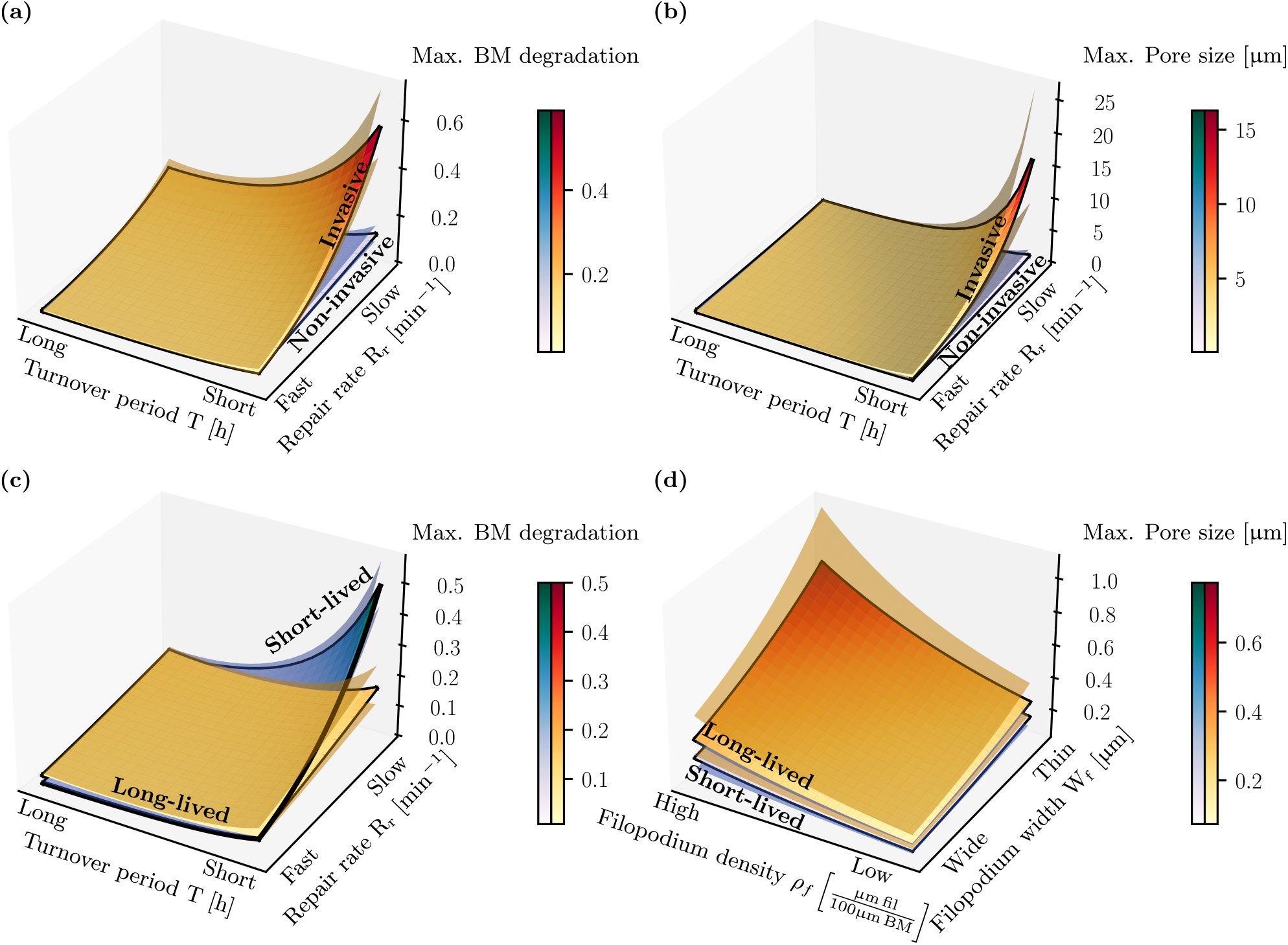
Sensitivity analysis of invasion dynamics on the regulation of CCLs and filopodial activity. Response surfaces were constructed by fitting a second-order metamodel of small spheroids (16 µm-diameter) during 48h, varying turnover period (T) from 2 to 17h, repair rate (R_r_) from 0.03125 to 0.5 per minute, density of filopodia (*ρ* _*f*_) from 20 to 50µm of filopodia per 100µm of BM, filopodium width (W_f_) from 0.5 to 2µm, and filopodium lifespan (L_f_) from 24 to 96 min. The fitted metamodels are defined in Eq. (19) and Table 4. The *z* axes represent response variables: maximum BM degradation in **(a)** and **(c)** and maximum pore size in **(b)** and **(d)**. Each response surface is shown with confidence intervals plotted as translucent surfaces. **(a)**-**(b)** Long turnover periods and fast BM repair protect against degradation **(a)** and pore enlargement **(b)**. Invasive surfaces are obtained with abundant and thin filopodia, while non-invasive ones are the opposite. **(c)** Filopodium lifespan alters the influence of the repair rate. Fixing density and width, long-lived filopodia are slightly more invasive at high repair rates, while short-lived ones are more invasive at low repair rates. **(d)** Filopodium lifespan amplifies the effect of abundant thin filopodia. Fixing turnover period and repair rate, long-lived filopodia promote larger pore enlargement than short-lived ones.

The analysis indicates that BM degradation and pore size are largely dominated by BM dynamics. Long turnover periods and fast repair effectively protect the BM: CCL deactivation events are rare, and any inactive link is quickly reactivated after filopodium retraction. This trend is illustrated in Fig. 7**(a)** and **(b)**, where BM degradation and pore size are plotted as functions of turnover period and repair rate. Each plot contains two surfaces: the red-yellow one corresponds to invasive filopodial activity and the blue-purple one to non-invasive filopodial activity. High BM degradation and large pore sizes occur only on the invasive surface, in regions with short turnover periods and slow repair.

The analysis indicates that such invasive activity is obtained in spheroids with abundant, thin filopodia (see Table 4). This is illustrated in Fig. 7**(d)**; the largest pores arise under conditions of high filopodial density and small width. Thin filopodia are more effective than wide ones since they can penetrate the BM more easily, allowing them to extend further outside. Since long filopodia take more time to retract, the likelihood that additional filopodia will arrive before the first ones retract is increased.

Lifespan modulates BM protectiveness and filopodial invasiveness (Fig. 7**(c)** and **(d)**). Long-lived filopodia slightly reduce the protective effect of fast repair, since their extended presence allows the arrival of new filopodia before link reactivation. In contrast, under slow repair, short-lived filopodia drive higher BM degradation. Since they affect more sites within the same time window, they promote a more widespread damage which in this case is not rapidly reversed. Lifespan also amplifies the invasive effects of density and width (Fig. 7**(d)**). Long-lived filopodia maintain CCL inactivity long enough to coincide with turnover of neighbouring links. Consequently, pore enlargement is promoted.

These results identify BM turnover period and BM repair rate as the key parameters that can enable protease-independent invasion. Only under short turnover and slow repair do spheroids with abundant, thin, and long-lived filopodia exhibit invasive behaviour.

## 4. Discussion

In this study, we have proposed a novel hypothesis for protease-independent BM invasion. Traditionally, it has been thought that tumour cells need MMPs to breach the BM [1]. However, recent evidence shows that even under protease inhibition, tumour cells can still invade [8], indicating the presence of protease-independent mechanisms. This was difficult to reconcile since BMs comprise a covalently linked network of collagen IV fibres. Yet, Wisdom et al. [18] revealed that filopodial activity can enlarge pores in plastic ECMs, and Corinus et al. [17] showed that in 3D mammary cell spheroids invasive cells protrude abundant filopodia coinciding with the formation of large pores, suggesting that filopodia actively prepare BM breaching. Moreover, Wisdom et al. [19] showed that ECM plasticity can be regulated by modulating the amount of CCLs in reconstituted BMs [19]. In other words, regions lacking CCLs can be exploited by filopodia to drive pore enlargement. We extended these ideas by suggesting that such regions lacking CCLs arise naturally through collagen IV turnover, leaving a brief window of opportunity for filopodia to occupy the vacant space.

To test this hypothesis, we developed a biophysical GS-PDE modelling framework coupling a deformable TCC to a dynamic BM undergoing collagen IV turnover. All parameters were derived from experimental data on membrane mechanics, filopodial activity, and collagen IV turnover (Table 2). Our biophysical model is able to reproduce the formation of long filopodia (57.1µm), in the same order of magnitude as those observed *in vitro* (up to 80µm [17]). Moreover, the results show that under suitable conditions, random turnover and filopodia can synchronise, leading to progressive pore enlargement. The model predicts that pore enlargement results from the collaboration of several filopodia entering and leaving the same region of the BM at different times. Although our results cannot demonstrate that this mechanism occurs *in vivo*, they place turnover as a plausible contributor to protease-independent invasion.

A key model prediction is that collagen IV turnover can only be exploited if the BM is not immediately rebuilt after filopodium retraction. This dependency is captured by the rate of BM repair, which can be linked to the levels of expression of BM core components. Corinus et al. [17], Fayad et al. [29] showed that the downregulation of BM core components facilitates invasion in mammary cell spheroids. Similarly, Spaderna et al. [28] demonstrated that the expression of BM components is locally reduced at invasive regions in colorectal cancer. These results suggest that filopodia can efficiently exploit collagen IV turnover provided that BM expression is reduced. Nevertheless, experimental evidence is needed to assess how quickly BM structure is repaired after local disruption.

Regarding filopodial activity, our GS-PDE model predicts that invasion is favoured by abundant filopodia, consistent with experimental observations [17]. Nonetheless, while thin filopodia favoured invasion in our GS-PDE, Corinus et al. [17] reported wider filopodia in invasive mammary cell spheroids. This discrepancy reflects differences between *in silico* and *in vitro* filopodia. In our GS-PDE model, filopodium width depends on the size of the region where the force is applied and on pore size: even if the force is highly localised in thin filopodia, a large pore leads to a wider filopodium base due to membrane tension and bending, key geometric properties of the energetic cell membrane. Thus, while the tip remains thin, the base adapts to the pore size, as seen in Fig. 6**(a)**. In contrast, *in vitro* filopodia do not necessarily adapt their width to the pore size. This suggests that thin filopodia are more efficient at initiating pore formation, while subsequent widening is required to sustain pore enlargement.

Filopodium lifespan also plays a dual role. Our GS-PDE model indicates that short-lived filopodia are more efficient at driving BM degradation when BM repair is slow, whereas long-lived ones enhance pore enlargement. Corinus et al. [17] reported that mammary cell spheroids display mostly thin, short-lived filopodia, but that invasive spheroids contain an additional small population of long-lived filopodia that eventually stabilise and widen. Hence, these results suggest a collaboration between filopodium populations: short-lived filopodia drive global BM degradation and long-lived filopodia drive local pore enlargement.

In summary, our work proposes a mechanism for protease-independent BM invasion based on collagen IV turnover and filopodial activity. Stochastic turnover transiently deactivates CCLs, creating a window of opportunity for filopodia to slip into. Abundant, thin, and short-lived filopodia rapidly occupy these sites and block reactivation. If BM repair is slow, successive filopodia reinforce local degradation, and turnover of neighbouring CCLs enables pore enlargement. Pore enlargement is then sustained by long-lived filopodia that eventually widen. Thus, this protease-independent mechanism suggests the existence of two complementary groups of filopodia: one for global degradation and the other for local pore enlargement at invasion sites.

Finally, the current modelling framework can be improved to capture additional phenomena. For instance, protease-degradation can be added in the rate of change of *c*_*l*_ (Fig. 2**(b)**) as a negative term dependent on filopodium presence. This would allow capturing the collaborative interplay of the filopodia and protease degradation in driving invasion. Likewise, extending the model to three dimensions would capture more realistic BM geometries and filopodial interactions. Furthermore, modelling nuclei would allow the evaluation of the full cell translocation through BM pores, providing a more complete description of the invasion process.

## 5. Acknowledgements

This project has been supported by the Institut national de la santé et de la recherche médicale through project INSERM-MIC2024-LSP 296457. This work has also been supported by the French government, through the UCA-JEDI Investments in the Future project managed by the National Research Agency (ANR) with the reference number ANR-15-IDEX-01. This work (AM) was supported by the Canada Research Chair (Tier 1) in Theoretical and Computational Biology (CRC-2022-00147), the Natural Sciences and Engineering Research Council of Canada (NSERC), Discovery Grants Program (RGPIN-2023-05231), the British Columbia Knowledge Development Fund (BCKDF), Canada Foundation for Innovation – John R. Evans Leaders Fund – Partnerships (CFI-JELF), and the British Columbia Foundation for Non-Animal Research. The authors would like to thank Julia Dubreuil and Sophie Abelanet for contributing to the experimental data shown in Fig. 1. The authors gratefully acknowledge the IdEx program of Université Côte d’Azur for partially supporting the collaboration between the Jean Alexandre Dieudonné Laboratory and the University of British Columbia.

## References

[1] R. Sekiguchi, K. M. Yamada, Basement Membranes in Development and Disease, Elsevier, 2018, pp. 143–191. doi:10.1016/bs.ctdb.2018.02.005.

[2] B. Alberts, A. Johnson, J. Lewis, D. Morgan, M. Raff, K. Roberts, P. Walter, Cell junctions, cell adhesion, and the extracellular matrix, in: Molecular Biology of the Cell, 6th ed., Garland Science, 2015.

[3] P. D. Yurchenco, Basement membranes: Cell scaffoldings and signaling platforms, Cold Spring Harbor Perspectives in Biology 3 (2010) a004911. doi:10.1101/cshperspect.a004911.

[4] N. Khalilgharibi, Y. Mao, To form and function: on the role of basement membrane mechanics in tissue development, homeostasis and disease, Open Biology 11 (2021). doi:10.1098/rsob.200360.

[5] G. A. Abrams, S. L. Goodman, P. F. Nealey, M. Franco, C. J. Murphy, Nanoscale topography of the basement membrane underlying the corneal epithelium of the rhesus macaque, Cell and Tissue Research 299 (2000) 39–46. doi:10.1007/s004410050004.

[6] A. Gaiko-Shcherbak, G. Fabris, G. Dreissen, R. Merkel, B. Hoffmann, E. Noetzel, The acinar cage: Basement membranes determine molecule exchange and mechanical stability of human breast cell acini, PLOS ONE 10 (2015) e0145174. doi:10.1371/journal.pone.0145174.

[7] K. Wolf, M. te Lindert, M. Krause, S. Alexander, J. te Riet, A. L. Willis, R. M. Hoffman, C. G. Figdor, S. J. Weiss, P. Friedl, Physical limits of cell migration: Control by ecm space and nuclear deformation and tuning by proteolysis and traction force, Journal of Cell Biology 201 (2013) 1069–1084. doi:10.1083/jcb.201210152.

[8] J. Chang, O. Chaudhuri, Beyond proteases: Basement membrane mechanics and cancer invasion, Journal of Cell Biology 218 (2019) 2456–2469. doi:10.1083/jcb.201903066.

[9] L. C. Kelley, Q. Chi, R. Cáceres, E. Hastie, A. J. Schindler, Y. Jiang, D. Q. Matus, J. Plastino, D. R. Sherwood, Adaptive f-actin polymerization and localized atp production drive basement membrane invasion in the absence of mmps, Developmental Cell 48 (2019) 313–328.e8. doi:10.1016/j.devcel.2018.12.018.

[10] K. A. Moore, T. Polte, S. Huang, B. Shi, E. Alsberg, M. E. Sunday, D. E. Ingber, Control of basement membrane remodeling and epithelial branching morphogenesis in embryonic lung by rho and cytoskeletal tension, Developmental Dynamics 232 (2004) 268–281. doi:10.1002/dvdy.20237.

[11] J. S. Harunaga, A. D. Doyle, K. M. Yamada, Local and global dynamics of the basement membrane during branching morphogenesis require protease activity and actomyosin contractility, Developmental Biology 394 (2014) 197–205. doi:10.1016/j.ydbio.2014.08.014.

[12] T. Shibue, M. W. Brooks, M. F. Inan, F. Reinhardt, R. A. Weinberg, The outgrowth of micrometastases is enabled by the formation of filopodium-like protrusions, Cancer Discovery 2 (2012) 706–721. doi:10.1158/2159-8290.cd-11-0239.

[13] G. Jacquemet, H. Hamidi, J. Ivaska, Filopodia in cell adhesion, 3d migration and cancer cell invasion, Current Opinion in Cell Biology 36 (2015) 23–31. doi:10.1016/j.ceb.2015.06.007.

[14] G. Jacquemet, H. Baghirov, M. Georgiadou, H. Sihto, E. Peuhu, P. Cettour-Janet, T. He, M. Perälä, P. Kronqvist, H. Joensuu, J. Ivaska, L-type calcium channels regulate filopodia stability and cancer cell invasion downstream of integrin signalling, Nature Communications 7 (2016). doi:10.1038/ncomms13297.

[15] G. Jacquemet, I. Paatero, A. F. Carisey, A. Padzik, J. S. Orange, H. Hamidi, J. Ivaska, Filoquant reveals increased filopodia density during breast cancer progression, Journal of Cell Biology 216 (2017) 3387–3403. doi:10.1083/jcb.201704045.

[16] R. Cáceres, N. Bojanala, L. C. Kelley, J. Dreier, J. Manzi, F. Di Federico, Q. Chi, T. Risler, I. Testa, D. R. Sherwood, J. Plastino, Forces drive basement membrane invasion in caenorhabditis elegans, Proceedings of the National Academy of Sciences 115 (2018) 11537–11542. doi:10.1073/pnas.1808760115.

[17] A. Corinus, S. Abelanet, J. Dubreuil, Z. Zhu, S. Pisano, C. Boscagli, A.-S. Gay, D. Debayle, M. Truchi, K. Lebrigand, S. Lacas-Gervais, F. Brau, X. Descombes, P. Rousselle, M. Franco, F. Luton, Human mammary 3d spheroid models uncover the role of filopodia in breaching the basement membrane to facilitate invasion, bioRxiv (2025). doi:10.1101/2025.08.29.672913.

[18] K. M. Wisdom, K. Adebowale, J. Chang, J. Y. Lee, S. Nam, R. Desai, N. S. Rossen, M. Rafat, R. B. West, L. Hodgson, O. Chaudhuri, Matrix mechanical plasticity regulates cancer cell migration through confining microenvironments, Nature Communications 9 (2018). doi:10.1038/s41467-018-06641-z.

[19] K. M. Wisdom, D. Indana, P.-E. Chou, R. Desai, T. Kim, O. Chaudhuri, Covalent cross-linking of basement membrane-like matrices physically restricts invasive protrusions in breast cancer cells, Matrix Biology 85-86 (2020) 94–111. doi:10.1016/j.matbio.2019.05.006.

[20] K. Tanner, Regulation of the basement membrane by epithelia generated forces, Physical Biology 9 (2012) 065003. doi:10.1088/1478-3975/9/6/065003.

[21] M. A. Morrissey, R. Jayadev, G. R. Miley, C. A. Blebea, Q. Chi, S. Ihara, D. R. Sherwood, Sparc promotes cell invasion in vivo by decreasing type iv collagen levels in the basement membrane, PLOS Genetics 12 (2016) e1005905. doi:10.1371/journal.pgen.1005905.

[22] D. P. Keeley, E. Hastie, R. Jayadev, L. C. Kelley, Q. Chi, S. G. Payne, J. L. Jeger, B. D. Hoffman, D. R. Sherwood, Comprehensive endogenous tagging of basement membrane components reveals dynamic movement within the matrix scaffolding, Developmental Cell 54 (2020) 60–74.e7. doi:10.1016/j.devcel.2020.05.022.

[23] S. Srinivasan, W. Ramos-Lewis, M. R. Morais, Q. Chi, A. W. Soh, E. Williams, R. Lennon, D. R. Sherwood, A collagen iv fluorophore knock-in toolkit reveals trimer diversity in c. elegans basement membranes, Journal of Cell Biology 224 (2025) e202412118. doi:10.1083/jcb.202412118.

[24] Y. Matsubayashi, B.J. Sánchez-Sánchez, S. Marcotti, E. Serna-Morales, A. Dragu, M.-d.-C. Díaz-de-la Loza, G. Vizcay-Barrena, R. A. Fleck, B. M. Stramer, Rapid homeostatic turnover of embryonic ecm during tissue morphogenesis, Developmental Cell 54 (2020) 33–42.e9. doi:10.1016/j.devcel.2020.06.005.

[25] R. A. Jones, B. Trejo, P. Sil, K. A. Little, H. A. Pasolli, B. Joyce, E. Posfai, D. Devenport, An mturq2-col4a1 mouse model allows for live visualization of mammalian basement membrane development, Journal of Cell Biology 223 (2023) e202309074. doi:10.1083/jcb.202309074.

[26] D. Wuergezhen, E. Gindroz, R. Morita, K. Hashimoto, T. Abe, H. Kiyonari, H. Fujiwara, An egfp-col4a2 mouse model reveals basement membrane dynamics underlying hair follicle morphogenesis, Journal of Cell Biology 224 (2024) e202404003. doi:10.1083/jcb.202404003.

[27] M. C. van den Berg, L. MacCarthy-Morrogh, D. Carter, J. Morris, I. Ribeiro Bravo, Y. Feng, P. Martin, Proteolytic and opportunistic breaching of the basement membrane zone by immune cells during tumor initiation, Cell Reports 27 (2019) 2837–2846.e4. doi:10.1016/j.celrep.2019.05.029.

[28] S. Spaderna, O. Schmalhofer, F. Hlubek, G. Berx, A. Eger, S. Merkel, A. Jung, T. Kirchner, T. Brabletz, A transient, emt-linked loss of basement membranes indicates metastasis and poor survival in colorectal cancer, Gastroenterology 131 (2006) 830–840. doi:10.1053/j.gastro.2006.06.016.

[29] R. Fayad, M. V. Rojas, M. Partisani, P. Finetti, S. Dib, S. Abelanet, V. Virolle, A. Farina, O. Cabaud, M. Lopez, D. Birnbaum, F. Bertucci, M. Franco, F. Luton, Efa6b regulates a stop signal for collective invasion in breast cancer, Nature Communications 12 (2021). doi:10.1038/s41467-021-22522-4.

[30] K. M. Yamada, J. W. Collins, D. A. Cruz Walma, A. D. Doyle, S. G. Morales, J. Lu, K. Matsumoto, S. S. Nazari, R. Sekiguchi, Y. Shinsato, S. Wang, Extracellular matrix dynamics in cell migration, invasion and tissue morphogenesis, International Journal of Experimental Pathology 100 (2019) 144–152. doi:10.1111/iep.12329.

[31] K.-A. Norton, M. Wininger, G. Bhanot, S. Ganesan, N. Barnard, T. Shinbrot, A 2d mechanistic model of breast ductal carcinoma in situ (dcis) morphology and progression, Journal of Theoretical Biology 263 (2010) 393–406. doi:10.1016/j.jtbi.2009.11.024.

[32] P. Macklin, M. E. Edgerton, A. M. Thompson, V. Cristini, Patient-calibrated agent-based modelling of ductal carcinoma in situ (dcis): From microscopic measurements to macroscopic predictions of clinical progression, Journal of Theoretical Biology 301 (2012) 122–140. doi:10.1016/j.jtbi.2012.02.002.

[33] G. D’Antonio, P. Macklin, L. Preziosi, An agent-based model for elasto-plastic mechanical interactions between cells, basement membrane and extracellular matrix, Mathematical Biosciences and Engineering 10 (2013) 75–101. doi:10.3934/mbe.2013.10.75.

[34] Y. Kim, M. A. Stolarska, H. G. Othmer, The role of the microenvironment in tumor growth and invasion, Progress in Biophysics and Molecular Biology 106 (2011) 353–379. doi:10.1016/j.pbiomolbio.2011.06.006.

[35] C. Shi, C.-H. Huang, P. N. Devreotes, P. A. Iglesias, Interaction of motility, directional sensing, and polarity modules recreates the behaviors of chemotaxing cells, PLoS Computational Biology 9 (2013) e1003122. doi:10.1371/journal.pcbi.1003122.

[36] A. Moure, H. Gomez, Phase-field model of cellular migration: Three-dimensional simulations in fibrous networks, Computer Methods in Applied Mechanics and Engineering 320 (2017) 162–197. doi:10.1016/j.cma.2017.03.025.

[37] Moure, H. Gomez, Phase-field modeling of individual and collective cell migration, Archives of Computational Methods in Engineering 28 (2019) 311–344. doi:10.1007/s11831-019-09377-1.

[38] Y. Wu, C. Qin, H. Xing, D. Sun, Lattice boltzmann modeling of individual and collective cell dynamics in the presence of fluid flows, Physics of Fluids 36 (2024). doi:10.1063/5.0231067.

[39] R. Allena, D. Aubry, ‘run-and-tumble’ or ‘look-and-run’? a mechanical model to explore the behavior of a migrating amoeboid cell, Journal of Theoretical Biology 306 (2012) 15–31. doi:10.1016/j.jtbi.2012.03.041.

[40] S. Deveraux, R. Allena, D. Aubry, A numerical model suggests the interplay between nuclear plasticity and stiffness during a perfusion assay, Journal of Theoretical Biology 435 (2017) 62–77. doi:10.1016/j.jtbi.2017.09.007.

[41] D. Cusseddu, L. Edelstein-Keshet, J. Mackenzie, S. Portet, A. Madzvamuse, A coupled bulk-surface model for cell polarisation, Journal of Theoretical Biology 481 (2019) 119–135. doi:10.1016/j.jtbi.2018.09.008.

[42] D. Hernandez-Aristizabal, D.-A. Garzon-Alvarado, C.-A. Duque-Daza, A. Madzvamuse, A bulk-surface mechanobiochemical modelling approach for single cell migration in two-space dimensions, Journal of Theoretical Biology 595 (2024) 111966. doi:10.1016/j.jtbi.2024.111966.

[43] X. Cao, E. Moeendarbary, P. Isermann, P. M. Davidson, X. Wang, M. B. Chen, A. K. Burkart, J. Lammerding, R. D. Kamm, V. B. Shenoy, A chemomechanical model for nuclear morphology and stresses during cell transendothelial migration, Biophysical Journal 111 (2016) 1541–1552. doi:10.1016/j.bpj.2016.08.011.

[44] Stinner, T. Bretschneider, Mathematical modelling in cell migration: tackling biochemistry in changing geometries, Biochemical Society Transactions 48 (2020) 419–428. doi:10.1042/bst20190311.

[45] M. Elliott, B. Stinner, C. Venkataraman, Modelling cell motility and chemotaxis with evolving surface finite elements, Journal of The Royal Society Interface 9 (2012) 3027–3044. doi:10.1098/rsif.2012.0276.

[46] L. S. Ryder, Y. F. Dagdas, M. J. Kershaw, C. Venkataraman, A. Madzvamuse, X. Yan, N. Cruz-Mireles, D. M. Soanes, M. Oses-Ruiz, V. Styles, J. Sklenar, F. L. H. Menke, N. J. Talbot, A sensor kinase controls turgor-driven plant infection by the rice blast fungus, Nature 574 (2019) 423–427. doi:10.1038/s41586-019-1637-x.

[47] M. Deserno, Fluid lipid membranes: From differential geometry to curvature stresses, Chemistry and Physics of Lipids 185 (2015) 11–45. doi:10.1016/j.chemphyslip.2014.05.001.

[48] F. Frey, T. Idema, More than just a barrier: using physical models to couple membrane shape to cell function, Soft Matter 17 (2021) 3533–3549. doi:10.1039/d0sm01758b.

[49] Shao, W.-J. Rappel, H. Levine, Computational model for cell morphodynamics, Physical Review Letters 105 (2010). doi:10.1103/physrevlett.105.108104.

[50] M. P. Neilson, J. A. Mackenzie, S. D. Webb, R. H. Insall, Modeling cell movement and chemotaxis using pseudopod-based feedback, SIAM Journal on Scientific Computing 33 (2011) 1035–1057. doi:10.1137/100788938.

[51] C. M. Elliott, T. Ranner, Finite element analysis for a coupled bulk-surface partial differential equation, IMA Journal of Numerical Analysis 33 (2012) 377–402. doi:10.1093/imanum/drs022.

[52] M. Frittelli, A. Madzvamuse, I. Sgura, Bulk-surface virtual element method for systems of pdes in two-space dimensions, Numerische Mathematik 147 (2021) 305–348. doi:10.1007/s00211-020-01167-3.

[53] R. J. Ju, A. D. Falconer, C. J. Schmidt, M. A. Enriquez Martinez, K. M. Dean, R. P. Fiolka, D. P. Sester, M. Nobis, P. Timpson, A. J. Lomakin, G. Danuser, M. D. White, N. K. Haass, D. B. Oelz, S. J. Stehbens, Compression-dependent microtubule reinforcement enables cells to navigate confined environments, Nature Cell Biology 26 (2024) 1520–1534. doi:10.1038/s41556-024-01476-x.

[54] R. Barreira, C. M. Elliott, A. Madzvamuse, The surface finite element method for pattern formation on evolving biological surfaces, Journal of Mathematical Biology 63 (2011) 1095–1119. doi:10.1007/s00285-011-0401-0.

[55] Y. Chen, J. S. Lowengrub, Tumor growth in complex, evolving microenvironmental geometries: A diffuse domain approach, Journal of Theoretical Biology 361 (2014) 14–30. doi:10.1016/j.jtbi.2014.06.024.

[56] Y. Chen, J. S. Lowengrub, Tumor growth and calcification in evolving microenvironmental geometries, Journal of Theoretical Biology 463 (2019) 138–154. doi:10.1016/j.jtbi.2018.12.006.

[57] J. W. Barrett, H. Garcke, R. Nürnberg, Parametric finite element approximations of curvature-driven interface evolutions, Elsevier, 2020, pp. 275–423. doi:10.1016/bs.hna.2019.05.002.

[58] J. W. Barrett, H. Garcke, R. Nürnberg, Parametric approximation of willmore flow and related geometric evolution equations, SIAM Journal on Scientific Computing 31 (2008) 225–253. doi:10.1137/070700231.

[59] T. C. Blake, J. L. Gallop, Filopodia in vitro and in vivo, Annual Review of Cell and Developmental Biology 39 (2023) 307–329. doi:10.1146/annurev-cellbio-020223-025210.

[60] Fischer-Friedrich, A. A. Hyman, F. Jülicher, D. J. Müller, J. Helenius, Quantification of surface tension and internal pressure generated by single mitotic cells, Scientific Reports 4 (2014). doi:10.1038/srep06213.

[61] S. Himbert, A. D’Alessandro, S. M. Qadri, M. J. Majcher, T. Hoare, W. P. Sheffield, M. Nagao, J. F. Nagle, M. C. Rheinstädter, The bending rigidity of the red blood cell cytoplasmic membrane, PLOS ONE 17 (2022) e0269619. doi:10.1371/journal.pone.0269619.

[62] S. Latha, G. Sivaranjani, D. Dhanasekaran, Response surface methodology: A non-conventional statistical tool to maximize the throughput of streptomyces species biomass and their bioactive metabolites, Critical Reviews in Microbiology 43 (2017) 567–582. doi:10.1080/1040841x.2016.1271308.

[63] B. Singh, R. Kumar, N. Ahuja, Optimizing drug delivery systems using systematic “design of experiments”. part i: Fundamental aspects, Critical Reviews in Therapeutic Drug Carrier Systems 22 (2005) 27–105. doi:10.1615/critrevtherdrugcarriersyst.v22.i1.20.

[64] K. L. Knight, Study/experimental/research design: Much more than statistics, Journal of Athletic Training 45 (2010) 98–100. doi:10.4085/1062-6050-45.1.98.

[65] Dziuk, Computational parametric willmore flow, Numerische Mathematik 111 (2008) 55–80. doi:10.1007/s00211-008-0179-1.

[66] J. W. Barrett, H. Garcke, R. Nürnberg, The approximation of planar curve evolutions by stable fully implicit finite element schemes that equidistribute, Numerical Methods for Partial Differential Equations 27 (2010) 1–30. doi:10.1002/num.20637.

[67] J. W. Barrett, H. Garcke, R. Nürnberg, Computational parametric willmore flow with spontaneous curvature and area difference elasticity effects, SIAM Journal on Numerical Analysis 54 (2016) 1732–1762. doi:10.1137/16m1065379.

[68] W. Huang, R. D. Russell, Adaptive Mesh Movement in 1D, Springer New York, 2010, pp. 27–135. doi:10.1007/978-1-4419-7916-2\_2.

[69] M. S. Alnæs, A. Logg, K.B. Ølgaard, M. E. Rognes, G. N. Wells, Unified form language: A domain-specific language for weak formulations of partial differential equations, ACM Transactions on Mathematical Software 40 (2014) 1–37. doi:10.1145/2566630.

[70] M. W. Scroggs, I. A. Baratta, C. N. Richardson, G. N. Wells, Basix: a runtime finite element basis evaluation library, Journal of Open Source Software 7 (2022) 3982. doi:10.21105/joss.03982.

[71] A. Baratta, J. P. Dean, J. S. Dokken, M. Habera, J. S. Hale, C. N. Richardson, M. E. Rognes, M. W. Scroggs, N. Sime, G. N. Wells, Dolfinx: The next generation fenics problem solving environment, 2023. doi:10.5281/ZENODO.10447666.

[72] C. de Boor, Good approximation by splines with variable knot, in: Numerical Solution of Differential Equations, Lecture Notes in Math. 363, Springer, Dundee, 1973, pp. 12–20.

[73] A. Mackenzie, M. Nolan, C. F. Rowlatt, R. H. Insall, An adaptive moving mesh method for forced curve shortening flow, SIAM Journal on Scientific Computing 41 (2019) A1170–A1200. doi:10.1137/18m1211969.

